# Mapping the Promiscuous Binding Interface of HOX-A11 with KIX by Experimentally Guided *in-silico* docking

**DOI:** 10.1101/2021.01.11.426190

**Authors:** Soumya Ganguly, Günter P. Wagner, Jens Meiler

**Affiliations:** Department of Chemistry, Center for Structural Biology, Vanderbilt University, TN 37232, USA; Yale Systems Biology Institute and Department of Ecology and Evolutionary Biology, Yale University, New Haven, CT, 06511 USA; Departments of Chemistry, Pharmacology and Biomedical Informatics; Center for Structural Biology and Institute of Chemical Biology, Vanderbilt University, TN 37232, USA

## Abstract

Transcription factors (TFs) regulate levels of transcription through a complex array of protein-protein interactions, thereby controlling key physiological processes such as development, stress response and cell growth. The transcription factor HOXA11 contains an intrinsically disordered regions (IDR) through which it interacts with CREB binding protein (CBP) and regulates endometrial development and function in eutherian mammals. The interaction between the IDR of HOXA11 and CBP was analyzed using computational docking guided by experimental constraints. HOXA11 IDR interacts with the KIX domain of CBP at two discrete sites – MLL and cMyb, mediated by sticky hydrophobic grooves on the surface of KIX. A five residue motif FDQFF on HOXA11 can interact both at cMyb and MLL site of KIX resulting in a promiscuous binding.

**Author Summary:** We demonstrate how the intrinsically disordered region (IDR) of transcription factor HOXA11 interacts at two distinct sites of the transcription coactivator CREB binding protein (CBP). By combining computational docking with limited experimental data we construct models of the complex of the KIX domain within CBP and a short helical segment within the IDR of HOXA11. The interaction between HOXA11 and CBP is believed to trigger the downstream expression of genes important in embryonic development.

## Introduction

Eukaryotic transcription factors (TFs) are dominated by proteins with intrinsically disordered regions (IDRs) [1, 2]. Many of these TFs consist of a well-structured domain and at least one IDR with low sequence complexity. Earlier studies of TFs have primarily focused on the role of the structured domain in transcription regulation. Recent studies have focused on the flexibility, promiscuity, and plasticity of the IDRs which provide unique functional properties in transcriptional regulation through protein-protein interaction. Developing mechanistic models to understand protein-protein interaction via IDRs has been challenging due to lack of information regarding their specific interacting partners under physiological conditions. Flexibility of the IDR and low binding affinity has made it often difficult to design experiments to gain structural insights.

HOXA11 is a TF key to morphogenesis of nearly every bilateral organisms [3]. In placental mammals (eutherians), physical interaction between HOXA11 and the TF FOXO1A has been implicated in regulating gene expression in endometrial stromal cells during pregnancy [4–6]. Full length HOX protein folds into a bundle of three alpha helices known as the homeodomain (HD) at the C-terminus while the N-terminus remains unstructured. The mechanism of transcriptional regulation by HOX proteins is thought to be mediated by interaction of a conserved C-terminal DNA binding HD with an A-T rich DNA target sequence [7, 8]. Our previous study showed that truncation of the conserved residues at the disordered N-terminal region of HOXA11 affects downstream expression of reporter genes [9]. In the same study, we identified the histone acetyltransferase CBP as a cofactor of HOXA11. In the absence of FOXO1A, HOXA11 is an intrinsic repressor [10]. In presence of FOXO1A, transactivation of HOXA11 is mediated by interaction of the disordered region with KIX domain of CBP [9]. We identified the residues FDQFF in the disordered region of HOXA11 as the KIX binding domain (KBD).

In our present study, we have explored the mode of interaction between the KBD of HOXA11 and KIX at atomic details. Experimental data from NMR titration studies is used to guide rigid body docking of KBD and KIX.

## Results

We used RosettaDock [11] to investigate the protein-protein interaction between KIX and KBD of HOXA11. Using Monte Carlo Minimization (MCM) steps, RosettaDock searches the conformational space of two interacting rigid bodies and their side-chains to determine the lowest free energy arrangement of docked bodies. In order to reduce the degrees of freedom involved, we docked only the KBD including a few flanking residues of HOXA11 instead of the entire N-terminal region of HOXA11 comprising of a 150 residues IDR (Figure 1). The KBD, identified from experimental data comprises of FDQFF at position 142-146 in the human protein consistent with the φ-x-x-−φ-−φ motif where φ is represented by a bulky hydrophobic residue. The five-residue φ-x-x-−φ-−φ motif is a well characterized KIX recognition domain observed in other proteins and peptides interacting with KIX [12–16]. For our docking study we decided to include 16 residues with five residues flanking the N-terminal region of KBD and six residues at the C-terminal portion. We denote these 16 residues as the extended KBD (eKBD). This region was chosen as it is predicted to form an α-helix (see below).

**Figure 1.**
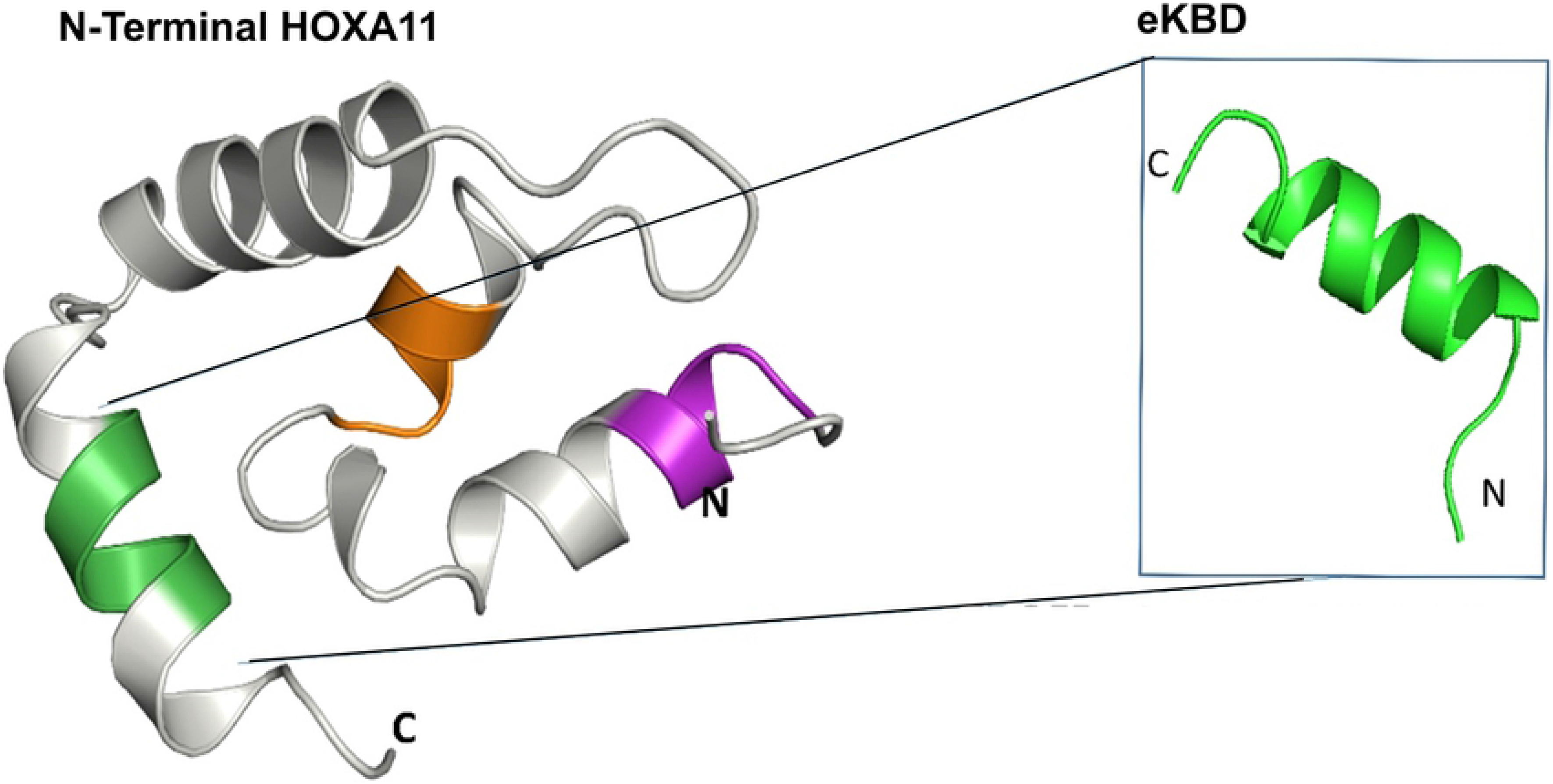
Lowest scoring *de novo* model of human N-terminal HOXA11 (residues 82-152). Zoomed in region is the 16 residue extended KIX binding domain (eKBD) consisting of FDQFF motif.

The *de novo* protein structure prediction algorithm, Rosetta [17] predicts that the N-terminal IDR of HOXA11 (residues 1-150) forms a loose helical bundle [9]. Depending on the starting and ending sequences this bundle consists of 3 to 4 α-helices. The 16 residue eKBD consistently assumes a helical conformation in the lowest energy models of N-terminal HOXA11. Energy minimized 3D helical structure of eKBD obtained from the lowest energy Rosetta model was used for rigid body docking in our present study. The three dimensional starting structure of KIX was derived from a previously determined NMR structure (PDB i.d. 2AGH) [13] after Rosetta energy minimization.

### Docking of MLL and cMyb peptide to KIX

To ensure the validity of our protocol, we benchmarked our docking process with MLL and cMyb peptides by individually docking them at their known binding sites. Experimentally determined models of these complexes exist [13, 18, 19]. Of the 39 MLL peptide residues only eleven residues have well defined α-helical structure. This eleven residue fragment of MLL was used for docking to KIX. We used seven previously identified residues of KIX demonstrating chemical shift perturbation in NMR studies upon MLL binding as atomic constrains to guide rigid body docking [19]. In the twenty lowest energy conformations generated during docking runs MLL occupies the KIX binding groove with a small difference in backbone rmsd (<0.4Å) to the experimentally determined conformation (Figure 2). The difference in backbone rmsd between the lowest energy docking model and the experiment was less than 0.3 Å. The minor difference between the experimental and docked model can be attributed to a slight rotation along the helical axis of the MLL peptide.

**Figure 2.**
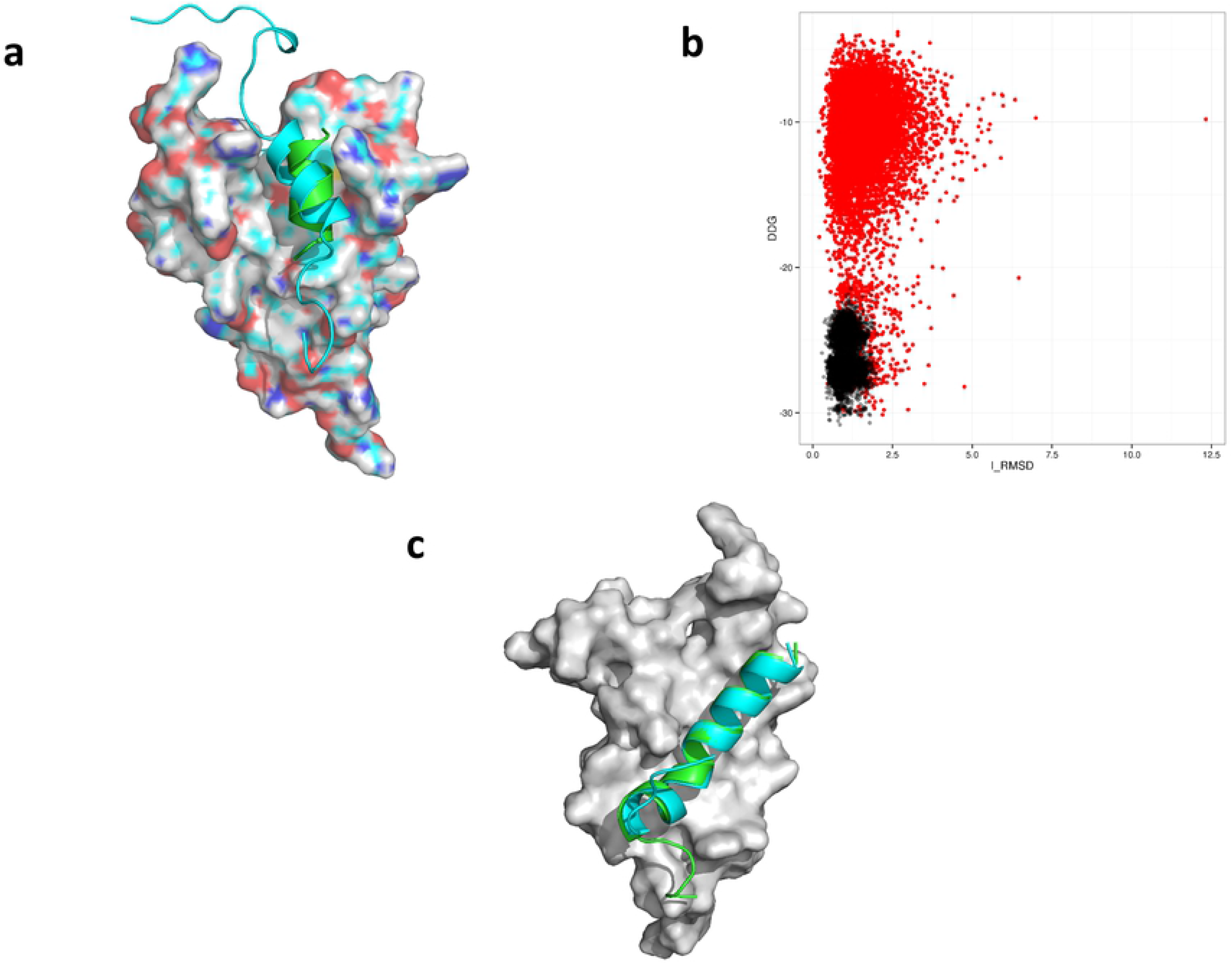
Controlled rigid body docking of MLL and cMyb peptide on KIX domain. (A) Overlay of the docked MLL peptide peptides from NMR structure in cyan and in green is the lowest ddg scoring model generated by RosettaDock. (B) A scattered plot of ddg vs i_rmsd from first (red) and second (black) docking runs of models generated when cMyb peptide was docked at the cMyb site of KIX. (C) Overlay of the docked cMyb peptide peptides from NMR structure in cyan and in green is the lowest ddg scoring model generated by RosettaDock.

Repeating the control docking run with the cMyb peptide also resulted in good agreement with experimental data. In this case all 24 resides of the cMyb peptide were used for docking. The hydrophobic groove where cMyb binds consists of L603, K606, L607, Y650, L653, A654 and I657 [18]. These seven residues were used as atomic constrains for docking of cMyb peptide. Significant improvement in overall interface rmsd (i_rmsd) and ddg score was observed during a high-resolution energy refinement of the complex. During this step, the top 1000 low scoring models from the low resolution docking is refined with a smaller rotation and translational perturbation [11](Figure 2). Experimental and lowest scoring docked model of cMyb occupy the same shallow hydrophobic groove with backbone rmsd less than 1.0 Å (Figure 2). The peptides from two models also exhibited near-perfect alignment along their α-helical axis.

Docking of MLL and cMyb peptides to KIX using experimentally guided data establishes that our protocol can be used to understand the atomic details of peptide-KIX interactions.

### Docking of eKBD to the MLL site

NMR titration of KIX with residues 80-152 of HOXA11 results in chemical shift perturbation of seven residues F612, T614, L620, K621, M625, E626 and N627 corresponding to KIX MLL site [9]. These residues were used as atomic constrains during docking of eKBD at the MLL site. Lowest energy conformations of eKBD were found to occupy the hydrophobic groove corresponding to the MLL site of KIX. All 20 low energy conformations were in a tight family with pair-wise backbone rmsd values of less than 1.25 Å. Although, both MLL and eKBD occupy the same hydrophobic groove they are offset axially by 7.8Å (Figure 3). Careful analysis of the interaction at atomic detail suggests a role of the FDQFF motif in the eKBD in HOXA11-KIX interaction. F142 and F147 of KBD were observed in a π-π stacking interaction with Y631 and F612 of KIX, respectively, at the MLL site. Interestingly, a few electrostatic interactions were also observed between the two domains notably between eKBD Q140 - KIX K656 and also between eKBD E147 and KIX K667 respectively (Figure 3).

**Figure 3.**
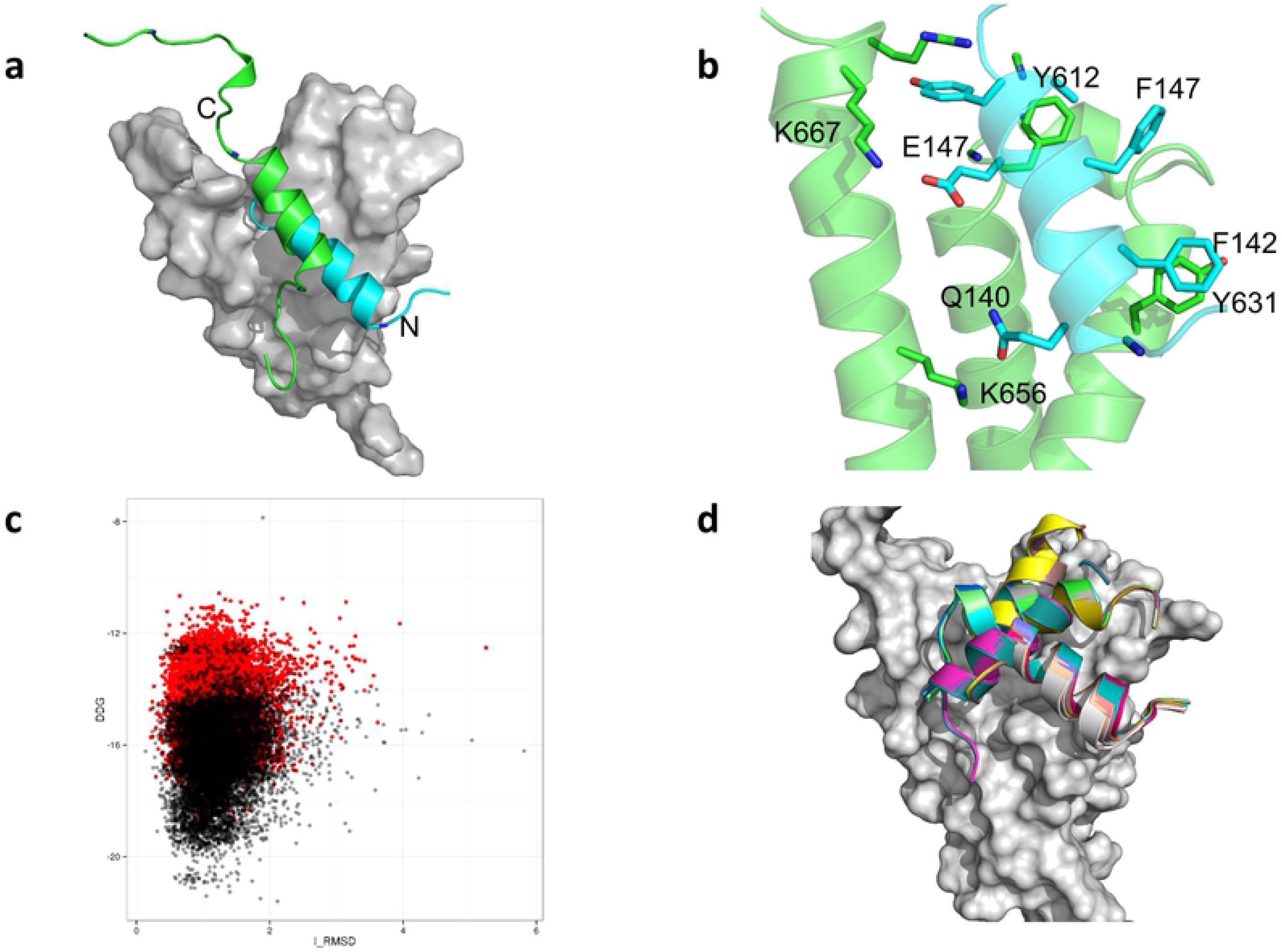
eKBD docking at MLL site of KIX. (A) An overlay of the lowest ddg docked model of eKBD (cyan) and MLL peptide from NMR structure (green) at the MLL site of KIX (grey). The N and C terminal orientation is similar for both peptides. (B) Details of the side-chain interactions between eKBD (cyan) and KIX (green). (C) Overlay of ddg vs i_rmsd score of 50,000 models from the refinement run for eKBD with FDQFF motif (black) and ADQAA (red). (D) Overlay of top 20 lowest scoring models from the docking of eKBD with ADQAA motif at the MLL site of KIX.

Mutation of phenylalanine to alanine at KBD of HOXA11 resulted in loss of chemical shift perturbation for KIX residues during NMR titration experiments [9]. We wanted to assess how this mutation affects the interaction of eKBD at the MLL site. Docking was repeated between eKBD and KIX where the FDQFF motif was replaced with ADQAA. As expected, the overall predicted free energy (ddg) of binding was significantly weaker for the ADQAA motif when compared with the FDQFF motif (Figure 3). Moreover, the top 20 low energy conformations from ADQAA docking were found to be more scattered across the MLL site with no direct alignment along the hydrophobic groove (Figure 3).

### Docking of eKBD at cMyb site

In the second experiment, we investigated the interaction between eKBD and KIX at the cMyb site. Two major conformations of eKBD were found to occupy the shallow hydrophobic groove at the cMyb site with an axial offset of 11 Å (Figure 4). In one conformation F145 was found to interact with Y650 of cMyb through π-π stacking (Figure 4). The primary residues mediating interaction between cMyb site and second lowest energy conformer are F146 – I657. Two more residues - V137 and L138 flanking the FDQFF motif were also involved in hydrophobic interaction with L603. We compared the overall i_rmsd vs ddg scores of eKBD from refinement dock runs at MLL and cMyb site. Interestingly, we found that the overall ddg score was lower for models docked at MLL than cMyb (Figure 4).

**Figure 4.**
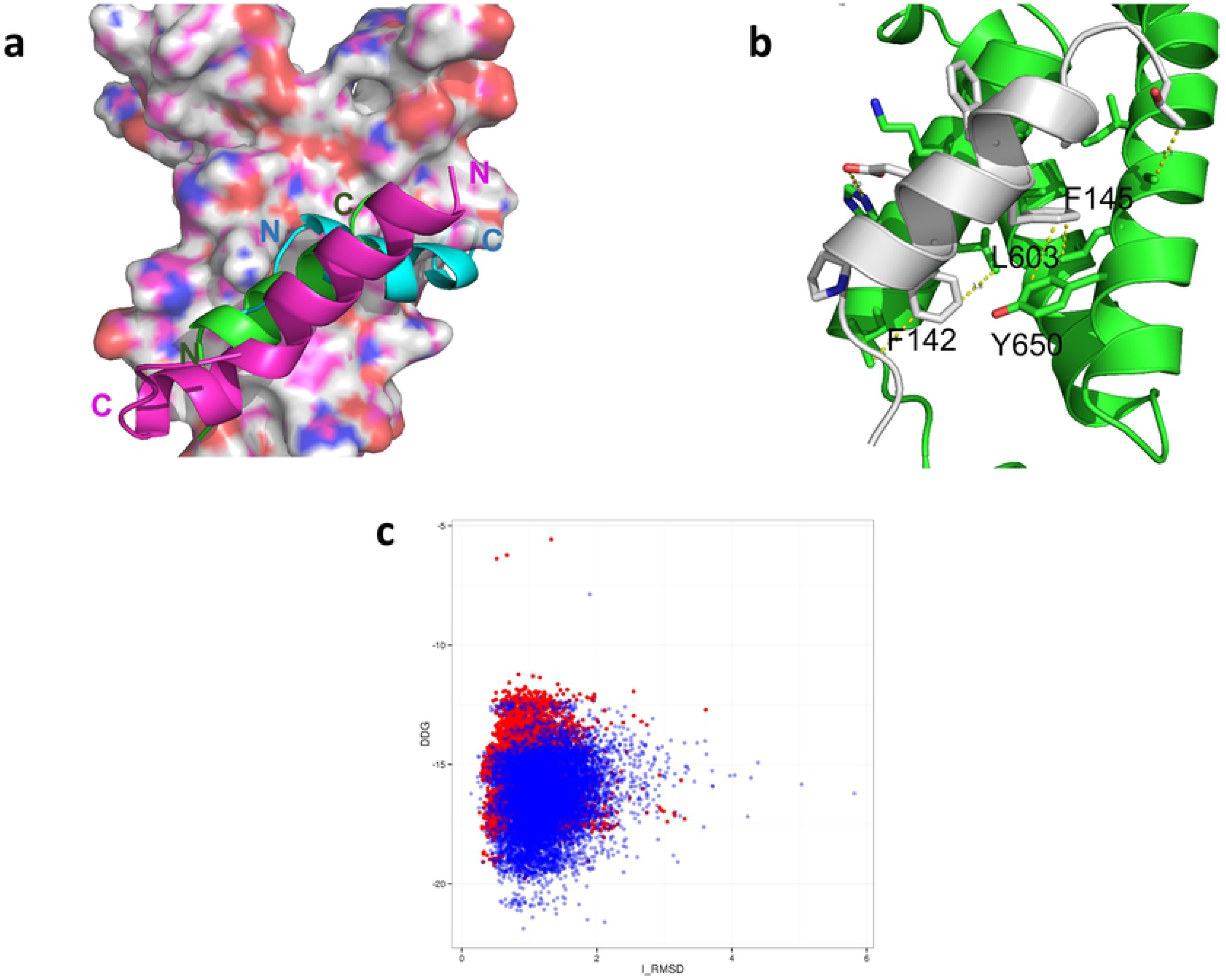
Docking of eKBD at cMyb site of KIX. (A) Overlay of two lowest energy (Green and Cyan) conformations of eKBD at the cMyb site of KIX. In purple is the docked model of cMyb peptide obtained from NMR structure. Note that the N-C terminal orientation of the cMyb peptide is oriented in opposite direction as compared to eKBD peptide. (B) Figure depicting the side chain interaction of one of the lowest energy conformer of eKBD with the cMyb site of KIX. (C) A scatter plot with an overlay of all 50,000 conformations from the refinement run of eKBD docked at MLL site (blue) and cMyb site (red).

## Discussion

Our previous study presented a mechanistic model of the functional interaction evolved between HOXA11 and FOXO1A. Intrinsic activation of HOXA11 was found to be regulated through intramolecular interaction between a CBP/P300 binding domain and a regulatory domain. The KIX domain of CBP was implicated in activation of HOXA11 through φ-x-x-φ-φ motif known as the KIX binding domain (KBD). NMR studies demonstrated that residues 142-146 of HOXA11 interacts at the MLL site and the cMyb site of KIX. Here we provide models at atomistic detail of the interaction using RosettaDock simulations guided by experimental data.

Docking of FDQFF results in an ensemble of low energy conformations occupying the MLL binding site with small deviation in backbone rmsd among the conformations. Moreover, mutation of the hydrophobic residues of FDQFF to alanine (ADQAA) results in conformations whose overall ddg score is higher than FDQFF suggesting lower affinity to the binding pocket. The top 20 low energy conformations of ADQAA display a larger variation in backbone rmsd. Many of the ADQAA low energy conformations did not completely occupy the MLL binding pocket. Our docking results suggest an increased affinity of FDQFF for the MLL binding site which diminishes upon mutation of the bulky hydrophobic residues. The docking results correlate well with experimental findings where similar mutation on the HOXA11 peptide show no interaction with KIX during NMR titration. The interaction between FDQFF and MLL site of KIX is dominated by hydrophobic residues of the two interacting partners.

The cMyb site of KIX is a long and shallow groove lined by side-chains of residues Leu603, Lys606, Leu607, Tyr650, Leu653, Ala654, Ile657 (JMB 2004). The top 20 low energy conformations of FDQFF were found in two unique conformations, both within the hydrophobic groove. In both conformations, the bulky phenylalanine residues were observed as principal driver of peptide protein interaction. The presence of a large number of hydrophobic sidechain on the cMyb site makes it ideal for FDQFF binding. FDQFF might assume several in-between conformation between the two low energy conformations shown here. Overall, the predicted binding free energy of docked models from cMyb site was higher than MLL suggesting weaker interaction at cMyb. Interestingly, we have observed similar weaker interaction at cMyb site in our earlier NMR titration of HOXA11 to KIX [9].

NMR titration and molecular docking study demonstrates promiscuity in binding with FDQFF interacting at two discrete surfaces of KIX. The binding affinity of FDQFF at the MLL and cMyb sites are also weak with a Kd in the range of hundreds of micro molar. Plasticity in interaction among binding partners can prove kinetically advantageous in recruiting the transcriptional cofactor to initiate complex formation as proposed by the fly casting model [20]. Promiscuous interaction coupled with low affinity binding can also help in fine tuning the transcriptional activity by adding spatial and temporal variability to the gene expression system.

Mechanism of transcriptional regulation involving HOXA11 is a dynamic process involving flexible domains along with conserved and well structure DNA binding domain. The flexible domain regulates gene expression through both inter and intramolecular interactions. Weak and promiscuous interaction also indicates subtlety involved in the transcription regulation perhaps through the formation of fuzzy complexes. Our studies were able to identify KIX as an important partner in the initiation of HOXA11 mediated transcription and the amino acid residues involved during the interaction.

## Methods

### Three Dimensional Structures of eKBD, KIX, MLL and cMyb

The 3-D structure of eKBD was obtained from a *de novo* folded N-terminal sequence of HOXA11 sequence generated by Rosetta [21]. Details of the *de novo* prediction of N-terminal HOXA11 has been discussed earlier [9]. From the *de novo* folded HOXA11 models, the lowest energy folded structure was identified as residues 82-152 of human HOXA11. This region assumes a loose helical conformation of which the 16 residue eKBD (GVLPQAFDQFFETAYGT) also assuming alpha helical structure. eKBD structure was isolated and relaxed using FastRelax protocol and used in docking studies.

Individual 3D- models of KIX, MLL and cMyb was obtained from the NMR structure of KIX bound to cMyb and MLL peptide (PDB ID 2AGH). From 20 low energy ensembles individual 3D structures were separated and relaxed. For the MLL peptide long unstructured N and C terminal residues were removed leaving only 11 residue helical fragment (SDIMDFVLKNT) for docking.

### Rigid Body Docking

Interaction between KIX and the peptide binding partners were investigated by RosettaDock. Since initial knowledge was available regarding the binding pocket, we performed local docking. Relaxed models of the docking partners are manually placed within ~10Å from the binding pocket before initiating docking protocol. During the first docking run 50,000 models were generated. For each individual docking step the peptide is randomly translated by 3Å and rotated by 8° in centroid model. During the refinement step the random perturbation was changed to 0.1 Å and 5°. Atomic constraints were used to guide binding partner into the pocket using experimental data. Residues of KIX exhibiting chemical shift perturbation upon peptide binding observed by NMR titration were used as constrains. Initial docking run generated 50,000 models which were analyzed based on ddg vs interface rmsd (irmsd) as compared to the lowest ddg structure. From this ensemble 1000 lowest scoring models were used as an input for a second docking run also called the refinement dock. During this run, coarse perturbation war set to 1 Å and 2.7° while refinement perturbations were 0.1 Å and 1°. The 50,000 models generated during refinement run was further analyzed based on their ddg score and interface rmsd. Finally, 20 lowest ddg models were identified and checked for consistencies in their docking pose.

## Acknowledgements

Financial support has been provided by a grant from the John Templeton Fund (JTF; grants #12793 and 54860). The opinions expressed in this paper are not those of the JTF.

